# Spatial contextual cueing is not a limiting factor for expert performance in the domain of team sports or action video game playing

**DOI:** 10.1101/267872

**Authors:** Anne Schmidt, Franziska Geringswald, Stefan Pollmann

**Author notes:** **Corresponding author:** Dipl.-Psych. Anne Schmidt, Otto-von-Guericke-Universität, Institut für Psychologie II, Postfach 4120, D-39016 Magdeburg, Germany.

## Abstract

We investigated in two experiments if handball and action video game players show improved implicit learning of repeated spatial configurations for efficient search guidance in comparison to a control group without sport or video game proficiency. To this end, we used both a sport-specific pseudo 3-D contextual cueing task and the original contextual cueing paradigm (Chun & Jiang, 1998). Contextual cueing was present in all groups. However, handball and action video game players did not differ in the strength of contextual cueing from the control group. Action video game players had shorter search times than controls in both experiments. In contrast, the handball players searched faster than controls in the sport-specific displays of Experiment 1 but not in the symbolic search task of Experiment 2. Thus, our findings provide no evidence that contextual cueing is a limiting factor for expert performance in the domain of team sports or action video game playing.

## Introduction

In a team sport like handball, players repeatedly encounter situations where a particular spatial constellation of players asks for a specific action. In these quickly changing situations, incidental learning of spatial contexts may occur that can optimize behavior in similar contexts.

In experimental psychology, such contextual cueing has even been observed in abstract visual search displays (Chun & Jiang; 1998). Spatial contexts can be learned incidentally, i.e. without the intention to learn, leading to more efficient visual search in repeated than in novel displays (reviewed by Chun, 2000; Goujon, Didierjean, & Thorpe, 2015). Both central and peripheral vision contribute to contextual cueing (Brady & Chun, 2007; Geringswald & Pollmann, 2015). Contextual cueing is flexible, it occurs even if only part of the target-distractor configuration is repeated (Brady & Chun, 2007; Song & Jiang, 2005) or if the trained display is rescaled, displaced or perceptually regrouped (Jiang & Wagner, 2004). Contextual cueing occurs rather automatically, even when attention is distracted away from the repeated items (Jiang & Leung, 2005) and when visuospatial working memory is loaded by a secondary task (Annac, Manginelli, Pollmann, Shi, Müller, & Geyer, 2013; Manginelli, Langer, Klose, & Pollmann, 2013; Vickery, Sussman, & Jiang, 2010). There is a debate about the implicit nature of contextual cueing (Geyer, Zehetleitner, & Müller, 2010; Smyth & Shanks, 2008; Vadillo, Konstantinidis, & Shanks, 2016). However, even if part of the repeated displays can be explicitly recognized, there is typically no correlation between explicit recognition scores and the size of the search facilitation in repeated displays. This suggests that contextual cueing is a genuine form of implicit learning (Colagiuri & Livesey, 2016).

Because of its incidental nature and its flexibility in utilizing even partial similarity in the structure of an environment, contextual cueing is likely to occur in team sport athletes and may be superiorly developed in elite athletes. However, to our knowledge, contextual cueing has not yet been investigated in athletes. Therefore, it is unknown if successful athletes, by training or selection (Kristjansson, 2013), may show enhanced benefits from contextual cueing.

Athletes have shown improved performance in a range of perceptual, attentional and cognitive tasks. To name just a few examples, improved basic perceptuomotor skills and visual search behavior have been reported (reviewed by Mann, Williams, Ward, & Janelle, 2007). Concerning attentional and cognitive skills, modulation of visuospatial attention was reported in elite volleyball players (Pesce-Anzeneder & Bösel, 1998). The nature of this modulation coincided with the demands of the game, e.g. more horizontal spread of attention was observed in soccer players versus more vertical spread in volleyball players (Hüttermann, Memmert, & Simons, 2014). Superior performance of athletes in comparison to non-athletes could also be demonstrated in a navigation task (Chaddock, Neider, Voss, Gaspar, & Kramer, 2011) and a multiple-object-tracking task (Faubert, 2013). Expert golf players showed superior performance in the Useful-Field-of-View task (Murphy, 2017). Conversely, training of perceptual functions improved several game parameters in professional baseball players (Deveau, Ozer, & Seitz, 2014). Generally, a meta-analytic review by Voss, Kramer, Basak, Prakash, & Roberts (2010) claimed superior performance for expert athletes on measures of processing speed and visual attention.

However, it is far from clear if elite athletes have improved attentional or cognitive skills that generalize to situations unrelated to sports. Moreover, athletes did not consistently show superior performance in attentional tasks. Memmert, Simons, & Grimme (2009) did neither find superior performance of expert handball players in multiple object tracking or the Useful-Field-of-View task nor less inattentional blindness. Expert basketball players did not show superior visuospatial working memory scores in the Corsi block tapping task (Furley & Memmert, 2010). Across many studies, sport-specific displays, stimuli, and processing requirements were more likely to lead to expert-novice differences (Abernethy, 1987b, 1988; Mann et al., 2007).

Contextual cueing in handball players was compared to two other groups, namely action video game players and controls without sport or video game proficiency. Action video game players were selected because enhanced attentional skills have been reported in this group, including improved visual control, greater attentional capacity, and better spatial allocation of attention (Green & Bavelier, 2003), enhanced target detection (Feng, Spence, & Pratt, 2007; Green & Bavelier, 2006a), and faster response selection (Castel, Pratt, & Drummond, 2005; Clark, Lanphear, & Riddick, 1987). For example, visual search performance and distractor inhibition was improved for action video game players relative to non-gamers (Bavelier, Green, Han, Renshaw, Merzenich, & Gentile, 2011; Buckley, Codina, Bhardwaj, & Pascalis, 2010; Chisholm, Hickey, Theeuwes, & Kingstone, 2010; Chisholm & Kingstone, 2012; Green & Bavelier, 2007; Hubert-Wallander, Green, Sugarman, & Bavelier, 2011). However, video game literature has come under severe criticism in recent years, mainly over poor research design, misinterpretation of data, and nonreplication of critical findings. Further issues (e.g. gender imbalance, different motivational levels, differential training histories of testing groups, lack of comparable groups) of video game studies are discussed by Kristjánsson (2013). Moreover, due to methodological deficits (e.g. inadequate control conditions) claims of video game advantages should be taken as tentative (Boot et al., 2011).

In a handball game as well as in an action video game, players have to respond quickly to changing spatial configurations. Clearly, playing handball additionally poses much more complex demands on appropriate motor actions. Given these commonalities and differences, we hypothesized that superior contextual cueing skills - i.e. memory-guided visual search in familiar scenes - might be beneficial for both kinds of players. Tentatively, we also hypothesized that handball players might have a greater training effect than action video game players because of the higher motor demands. Therefore, we reasoned that expert handball players, like expert action video game players, might demonstrate increased proficiency to learn repeating spatial contexts and use them for memory guided visual search.

In order to test this hypothesis, we investigated if team sport (handball) and action video game players show improved search guidance by repeated spatial configurations. In the first experiment, we used a search task designed to be advantageous for the handball players (search of the ball-carrying player in a playing field). In the second experiment, a typical symbolic contextual cueing paradigm (search of a “T” among “L”-shapes) was used in order to investigate the generality of a potential contextual cueing advantage for handball players and action video game players.

## Experiment 1

A general aim of the study was to investigate how the attentional skills of experts (team sport athletes, action video game players) compare with those of non-experts. Research on competitive sports (Mann et al., 2007; Voss et al., 2010) and video game playing (Boot et al., 2008; Green & Bavelier, 2003) indicates that attentional abilities may improve by practice and experience (Furley & Memmert, 2011). An example for such an attentional training approach may be the contextual cueing paradigm (Kristjánsson, 2013). The contextual cueing task may reflect attentional effects, elucidating if handball- and action video game playing improves context learning skills (see Faubert, 2013).

Chua and Chun (2003) emphasize the ecological validity of contextual cueing by arguing that contextual information are predominant in our environment, so that attention needs to be focused on relevant aspects and objects to allow effective action in the environment (Chun, 2000). However, the two-dimensional stimuli traditionally used in contextual cueing tasks limits the universality of contextual cueing (see Chua & Chun, 2003). Chua and Chun (2003) replicated the original contextual cueing task with volumetric pseudo-naturalistic 3-D shapes to represent the depth of the real world in order to improve ecological validity of contextual cueing. Results indicate that contextual cueing can be generalized to pseudo 3-D scenes. In order not to miss contextual cueing advantages that might be tied to handball-specific constellations, we started in Experiment 1 with a sport-specific pseudo 3-D contextual cueing task in which our participants had to search in displays that resembled players in a handball field. We reasoned that contextual cueing generalizes to sport-specific scenes with pseudo-3-D layouts and that subjects implicitly learn the spatial context in this novel contextual cueing task.

## Methods

### Participants

A total of 90 healthy participants (control: n=31, mean age = 24.8 years; handball players: n=31, mean age = 20.4 years; action video game players: n=28; mean age = 23.1 years) was recruited for this study. The athletes were subdivided into two categories, according to age and years of training: adult and junior. Nine adult players (9 women) from a 3rd league handball team and 22 junior players from the Sportclub Magdeburg (18 men and 4 women) playing in the A and B youth national league participated in this study. The junior handball players practice 6-7 days (15 hours) a week, whereas adult players have 3-5 days (8 hours) training per week. Junior handball players had more than 6 years and adult players more than 10 years handball experience. Controls (male = 10, female = 21) and action video game players (male = 23, female = 5) were recruited from the University of Magdeburg. Action video game players had to fulfill the following criteria: action video gamers needed to play action video games (e.g. Call of Duty, Battlefield) for a minimum of five hours a week for at least one year. Participants without any team sport and little to no action video game experience (less than 1 hour per week) were classified as controls.

Informed written permission was acquired prior to the experiments. Subjects were remunerated with course credits or received a payment of Euro 7. Further, subjects were naive about the purpose of the experiment. All participants had normal or corrected-to-normal vision. The experiments were approved by the Ethics Committee of the University of Magdeburg.

### Apparatus and Stimuli

The experiment was programmed and performed using the OpenGL-Psychophysics Toolbox extensions (Brainard, 1997; Pelli, 1997) in MATLAB (The MathWorks, Natick, MA, USA). Stimuli were displayed by a projector on a back-projection screen (1150 mm (1024 pixels) wide and 800 mm (768 pixels) high, vertical refresh rate of 60 Hz). Participants viewed the stimuli from a distance of approximately 126 cm (pixel size of 0.048° × 0.046°). Subjects completed the experiment individually in a dimly lit, sound-attenuated chamber.

The stimuli were generated with Blender 2.69 (Stichting Blender Foundation, Amsterdam, Netherlands). The texture of the stimuli was created with GNU Image Manipulation Program (GIMP 2.8). Each search display comprised one target (player with ball in left or right hand) and 11 distractors (players without ball), forming a spatial layout in pseudo 3-D space (Figure 1). The arms of the players could be positioned as follows: both arms up, both arms down, left up and right down, right up and left down. Stimuli subtended a minimum 1.3° × 1.9° and a maximum of 2.1° × 4.3° of visual angle. The ball subtended a minimum 0.2° × 0.2° and a maximum 0.5° × 0.5° of visual angle. The items were presented on a green pitch with a black background. All items were colored in brown (skin), white (shorts) and blue (shirt) to improve visibility. Stimuli were smaller the farther back they were to generate a sense of depth (Chua & Chun, 2003). Display size of the projected display can be seen in Figure 1.

**Figure 1.**
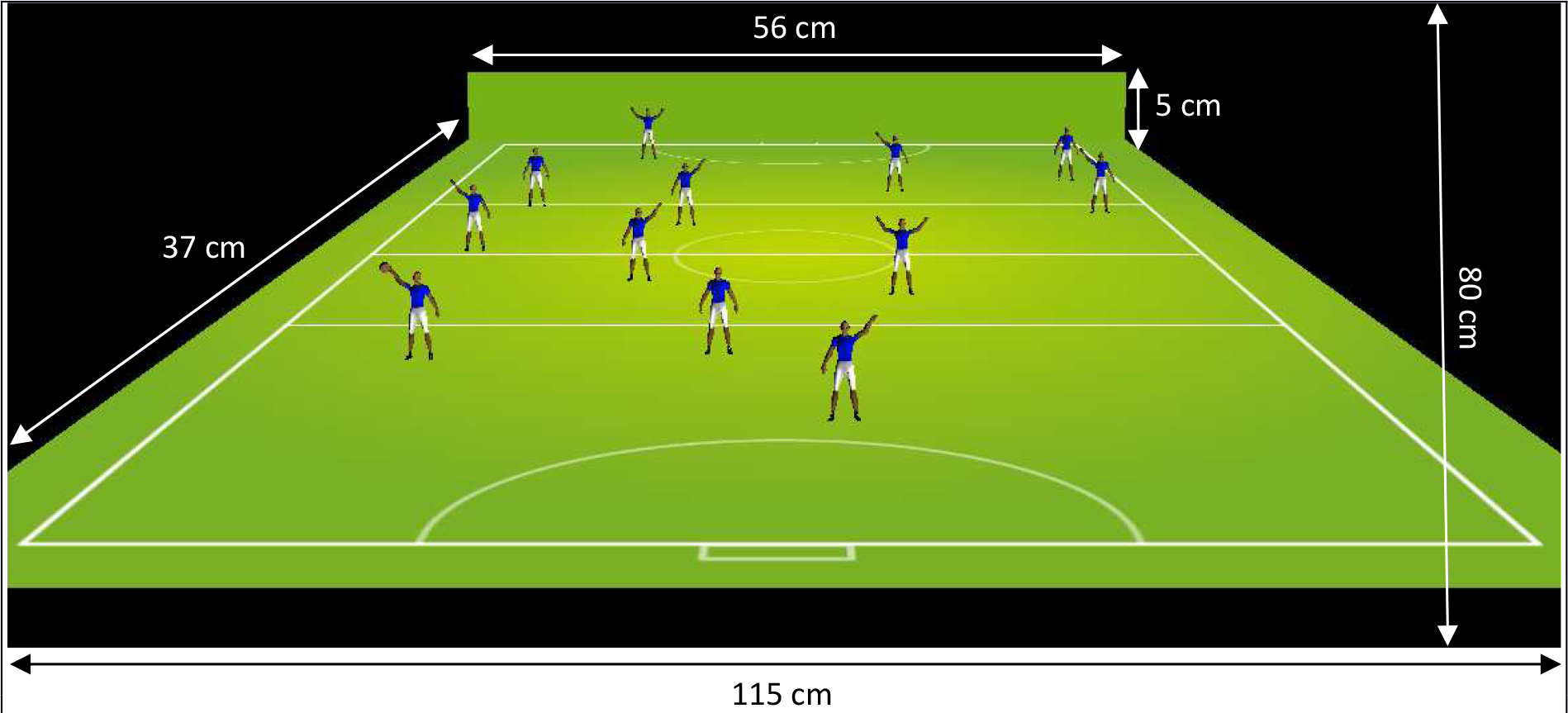
Display size of the projected display in Experiment 1.

Blocks contained 24 trials, 12 for each configuration type (repeated vs. novel). The repeated displays comprised 12 configurations that were repeated across blocks (1-20). 12 randomly chosen target positions were each paired with unique distractor configurations that remained constant throughout the experiment (repeated condition). The other 12 target positions were paired with distractor configurations that were newly generated for each block (novel condition). The targets in the novel sets appeared, as for the repeated configurations, equally often in one of 12 locations to control for target location repetition effects (Chun & Jiang, 1998). Stimuli were positioned in an invisible 3 column × 4 row grid, for a total of 12 rectangles in which objects could appear. The target positions and the eccentricity (near or far) were balanced across rectangles and conditions. The fixation stimulus was designed as a white cross at the center of the upper goal and subtending an area of 1.6° × 1.6°. The visual angle of the search display on the projection screen extended an area of 49.1° × 35.2°.

### Procedures

Participants searched for the player with the ball (target) among an array of distractor players and indicated the ball’s location (left vs. right hand) with left and right arrow keyboard button presses with the right hand. Each session started with a short training period to familiarize participants with the task, followed by the main search experiment and a recognition test. In the training phase, participants were shown 24 randomly generated displays which were not used in the main experiment. The testing phase comprised 20 blocks of 24 trials, 12 for each configuration type (repeated vs. novel displays). The entire experiment lasted approximately 35 minutes.

Each trial started with a blank interval for 500 ms followed by the fixation cross for 1000 ms. After a brief pause of 200 ms, the array of stimuli appeared on the projection screen (Figure 2). Participants were told to respond as quickly and accurately as possible. They were further instructed to search for the target passively (“be as receptive as possible and let the unique item ‘pop’ into your mind as you look at the screen”) as proposed by Lleras and von Mühlenen (2004). The search display remained on the screen until response. Auditory feedback was provided for correct (a 500-Hz low-pitch tone) and incorrect answers (a 1500-Hz high-pitch tone). At the end of the search task, the participants performed a recognition test, to evaluate whether repeated displays were explicitly remembered. The recognition test consisted of 24 trials, including the original 12 repeated and another 12 novel randomly generated configurations, presented in randomized order. Participants had to indicate by keyboard button press whether they had seen the displays during the course of the experiment or not. No feedback was given in the recognition task.

**Figure 2.**
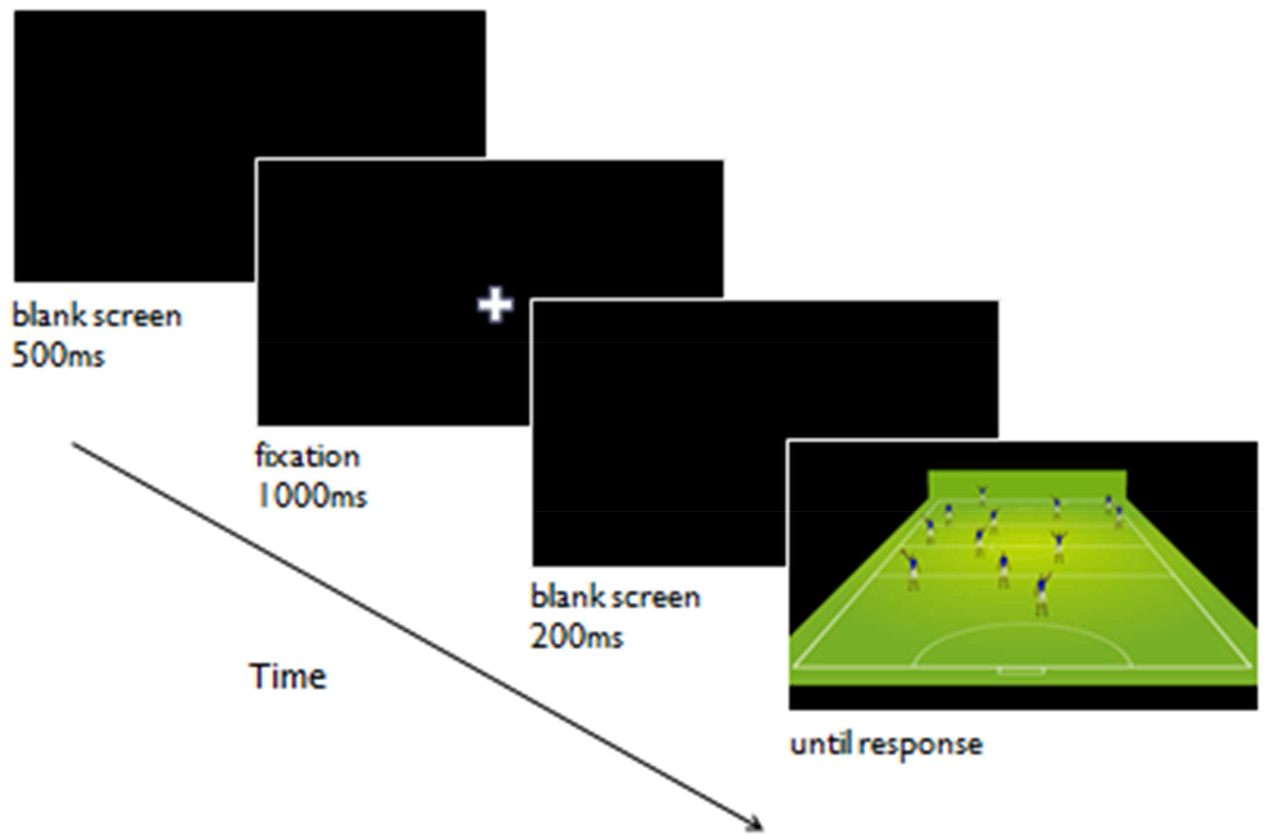
Procedure of a trial in Experiment 1. A trial consisted ofthe presentation of a blank screen [500ms], a fixation cross [1000ms], a blank screen [200ms], and the search display [presented until response].

### Data Exclusion Criteria

Two data exclusion criteria were applied for the search time data. First, all incorrect responses were removed from the data set. Second, trials in which the search time was shorter than 200 ms or larger than 3.5 standard deviations from the participants’ average search time in the remaining trials were discarded. The exclusion criteria led to the rejection of 3.4% (*SD* = 1.2%) of invalid data for the controls, 4.2% (*SD* = 1.8%) for handball players, and 4.4% (*SD* = 2.4%) for action video game players.

## Results

### Search Times

All statistical calculations were carried out using R-statistics (R Development Core Team, 2007). For the reaction time analysis, blocks were aggregated into four epochs, each containing five blocks. Averaged reaction times for the four epochs for repeated and novel displays separated by the three groups (control, handball, video) are displayed in Figure 3.

**Figure 3.**
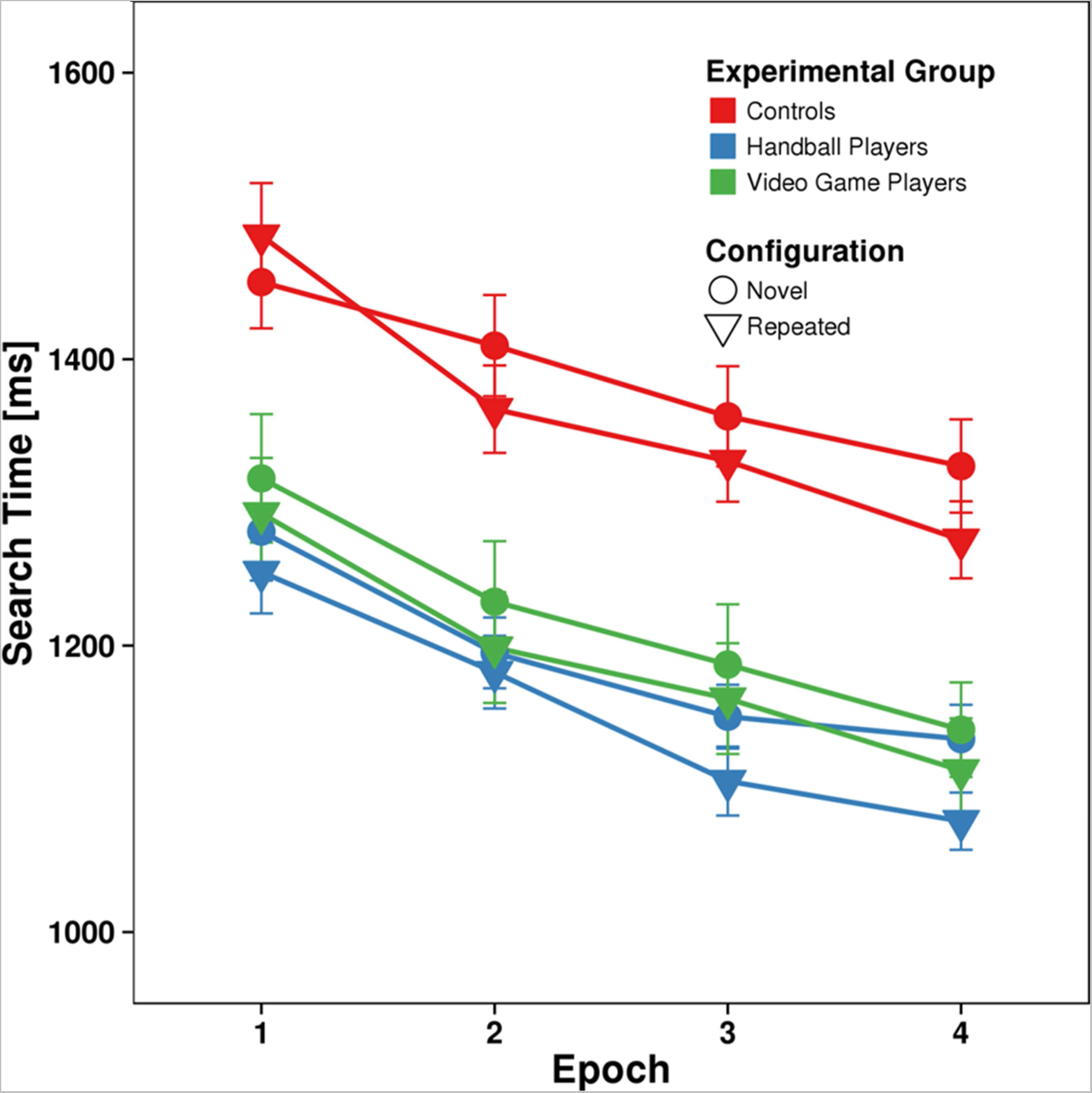
Search performance in Experiment 1. Averaged search times in the visual search task for the three groups controls (red), handball players (blue), and video game players (green) for repeated (triangles) and novel (circles) display as a function of epoch. Error bars represent the standard error of the mean.

A repeated-measures ANOVA with the within-subject factors configuration (repeated, novel) and epoch (1, 4) and the between-subject factor group (control, handball, video) was performed on mean reaction times. The first and the last epoch of the search experiment were contrasted to maximize effects due to learning.

The main effect of group was significant, *F*(2,87) = 13.757, *p* < 0.05. Post-hoc t-tests revealed that overall search speed was comparable between handball players (1186 ms) and action video game players (1216 ms), *t*(47) = 0.701, *p* = 0.49, but both groups outperformed controls (1385 ms, handball players: *t*(58) = 5.404, *p* < 0.05; action video players: *t*(53) = 3.687, *p* < 0.05). The significant main effect of epoch (*F*(1,87) = 247.014, *p* < 0.05) reflected general learning from the first (1348 ms) to the last epoch (1179 ms). In addition, we observed a significant main effect of configuration, *F*(1,87) = 6.248, *p* < 0.05, indicating that search in the repeated displays (1251 ms) was faster than search in novel displays (1277 ms). The significant interaction of epoch × configuration (*F*(1,87) = 6.997, *p* < 0.05) reflected increasingly faster search in repeated displays over time. While search times decreased by 149 ms in novel displays from the first to the last epoch, the use of contextual cues speeded up search by 189 ms in repeated displays. No other interaction effects were significant (all *F* < 2.57, *p* > 0.05) although the group × configuration × epoch interaction narrowly missed significance, *F* = 2.565, *p* = 0.08. Contextual cueing benefits in controls could potentially have been inflated by their overall slowed search compared to handball players and action video game players. To eliminate the effect of overall search speed, we performed an additional two-factorial mixed-design ANOVA with the within-subjects factor epoch and the between-subjects factor group on normalized contextual cueing effects. Normalized contextual cueing was obtained by dividing the absolute response time differences between novel and repeated displays by the response time of novel displays for each participant and each epoch. The ANOVA revealed a significant main effect of epoch, *F*(1,87) = 9.061, *p* < 0.05, confirming the increase of contextual cueing over time. Neither the main effect of group, *F*(2,87) = 0.909, *p* > 0.05, nor the interaction between epoch and group, *F*(2,87) = 1.239, *p* > 0.05, were significant.

### Age Differences

To investigate if the age of the handball players influenced the results, we ran a comparison of junior versus adult handball players. A repeated-measures ANOVA with the within-subject factors configuration (repeated, novel) and epoch (1, 4) and the between-subject factor group (junior handball players, adult handball players) was performed on mean reaction times. We observed significant main effects of epoch, F(1,29) = 70.967, p < 0.05, and configuration, F(1,29) = 5.864, p < 0.05. No group or interaction effects were significant (all F < 1.20, p > 0.05), indicating that junior and adult handball players did not differ in the contextual cueing task.

### Accuracy

Accuracies ranged from 96.7 to 100.0% (average 98.7%) in the control group, from 95.4 to 100.0% (average 98.3%) for the handball players and from 93.8 to 99.8% (average 98.3%) for the action video game players. An ANOVA on errors with the between-subject factor group (controls vs. handball players vs. videogame players) and the within-subject factor configuration (repeated vs. novel) yielded a significant main effect of configuration, *F*(1,87) = 10.976, *p* < 0.05, due to higher accuracies for repeated (average 98.6%) than for novel displays (average 98.2%). Neither the main effect of group, *F*(2,87) = 1.056, *p* > 0.05, nor the interaction between group and configuration, *F*(2,87) = 0.593, *p* > 0.05, was significant.

### Recognition Task

The control group reached a mean recognition accuracy of 50.4% (*SD* = 9.2%). Controls correctly reported repeated displays as ‘repeated’ (hit rate) on 45.2% (*SD* = 13.6%) of trials and erroneously categorized 44.4% (*SD* = 13.7%) of novel displays as repeated (false alarm rate). Hit and false alarm rates were not significantly different, *t*(24) = 0.244, *p* > 0.05, giving no indication that control participants were able to recognize repeated displays. In order to explore the relationship between recognition accuracy and contextual cueing effect we measured correlations between the normalized contextual cueing effects of the last epoch of the experiment with recognition accuracy. In the control group the standardized contextual cueing effect did not significantly correlate with the recognition accuracy (*r* = −0.156, *p* > 0.05).

Mean recognition accuracy for handball players was 48.7% (*SD* = 13.5%) with a mean hit rate of 48.4% (*SD* = 19.4%) and a mean false alarm rate of 51.1% (*SD* = 16.8%). The difference between hits and false alarms was not significant, *t*(24) = 0.556, *p* > 0.05. The normalized magnitude of contextual cueing correlated negatively with the recognition accuracy (*r* = −0.275, *p* < 0.05).

Action video game players yielded a mean recognition accuracy of 49.3% (*SD* = 8.8%). The mean hit rate (52.1%, *SD* = 13.9%) was comparable to the mean false alarm rate (53.6%, *SD* = 14.1%). The comparison between hit and false alarm rates during recognition did not indicate that video game players recognized repeated arrays, *t*(24) = 0.448, *p* > 0.05. Recognition accuracy did not correlate with the normalized contextual cueing effect (*r* = −0.017, *p* > 0.05).

## Discussion

Mean search times in Experiment 1 were shorter for both experimental groups in comparison to the control group. Search became faster for repeated relative to novel displays during the course of the experiment, indicating contextual cueing of visual search by repeated target-distractor configurations. However, we did not find increased contextual cueing in either handball players or video game players compared with controls.

Both handball players and video game players, however, had faster search times for both repeated and novel displays from the beginning of the experiment. The amount of general reduction of search times for both novel and repeated displays in the course of the experiment did not differ between groups.

In sum, both team sport athletes and action video game players were faster in finding the target, but neither was better than the control group in using repeated spatial configurations for search guidance.

Our results show that contextual cueing generalizes to sport-specific scenes with pseudo-3-D layouts. Similar findings have been reported by Brockmole & Henderson (2006) with real-world scenes and by Chua and Chun (2003) with pseudo naturalistic three-dimensional search arrays.

Furthermore, we did not observe evidence for explicit learning of spatial context in this novel contextual cueing task. These findings are in line with previous research for two-dimensional (2-D) layouts (Chun & Jiang, 1998, 1999) and for three dimensional (3-D) volumetric shapes (Chua & Chun, 2003). However, the evidence in favor of implicit contextual cueing is only tentative, because the statistical power of the recognition test - limited by the number of repeated displays - was far lower than that of the contextual cueing experiment (Vadillo et al., 2016).

## Experiment 2

There is a current debate in the expertise literature if basic cognitive skills can be improved as a function of prolonged practice in a certain field of expertise (e.g. sports, video game playing) or whether expert performance is domain specific. According to the narrow transfer hypothesis (Simons & Chabris, 2010), experts in activities such as team sports, action video game playing, or chess merely differ in cognitive abilities associated with their own performance environment (e.g. anticipation of ball directions in sport) and improvements in basic cognitive skills (e.g. memory capacity, intelligence) are not likely. On the other hand, there is evidence that expert performers are able to improve on a general cognitive level due to extensive training (Voss et al., 2010; Vestberg, Gustafson, Maurex, Ingvar, & Petrovic, 2012). This broad transfer hypothesis suggests that prolonged experience enables expert performers to enhance in basic cognitive skills and that those skills are translatable to different domains. Indeed, expert performers acquire specific knowledge in order to cope with specific constraints of their field of expertise (Starkes & Ericsson, 2003; Mann et al., 2007). But the question, if experts improve on a general cognitive skill level remains unanswered.

While the display in Experiment 1 was designed to resemble a handball playing field in order to tap into a potential sport-specific contextual cueing advantage of the handball players, the goal of Experiment 2 was to investigate if either handball playing or action video game playing may generalize to a contextual cueing advantage in arbitrary environments. In order to test this hypothesis, we used a ‘T’ among ‘L’ shape search (Chun & Jiang, 1998) in order to investigate contextual cueing.

## Methods

### Participants

A total of 75 healthy subjects (control: n=25, mean age = 25.1 years; handball players: n=25, mean age = 20.2 years; action video game players: n=25; mean age = 23.2 years) participated in Experiment 2. Eleven adult players (6 men, 5 women) from a 3rd league handball team and 14 junior players (14 men) playing in the A and B youth national league were recruited for Experiment 2. Action video game players (male = 11, female = 14) and controls (male = 22, female = 3) met the same criteria as in Experiment 1. Ten participants already participated in Experiment 1 (3 handball players, 7 video game players). Subjects were again compensated with course credits or received a payment of Euro 7. All participants had normal or corrected-to-normal vision. Moreover, subjects were not informed about the purpose of the study until they had completed the testing. The experiment was approved by the Ethics Committee of the University of Magdeburg. Informed written permission was acquired prior to the experiments.

### Apparatus & Stimuli

The apparatus was the same as described in Experiment 1. Search displays contained one target (90° or 270° rotated T) and 11 distractors (0°, 90°, 180°, 270° rotated L) with each item subtending 1.9° × 1.9° (Figure 4). An offset of 0.14° between the two segments of the L-shapes was chosen to increase search difficulty. The orientation of the target and the orientation of distractors were randomly chosen for each trial. A black cross (2.4° × 2.4°) at the center of the display was used as a fixation stimulus. Stimuli were black displayed on a gray background. The items were randomly positioned on four non-visible concentric circles with radii of 4°, 8°, 12°, and 16° visual angle each corresponding to 4, 12, 20, and 28 equidistant possible item locations. As in Experiment 1, blocks comprised 24 trials, 12 for each configuration type (repeated vs. novel). The positions of the items were balanced across quadrants (i.e., each quadrant of the display always comprise three search items) and configuration type. The visual angle of the search display on the projection screen extended an area of 49.8° × 37.3°.

**Figure 4.**
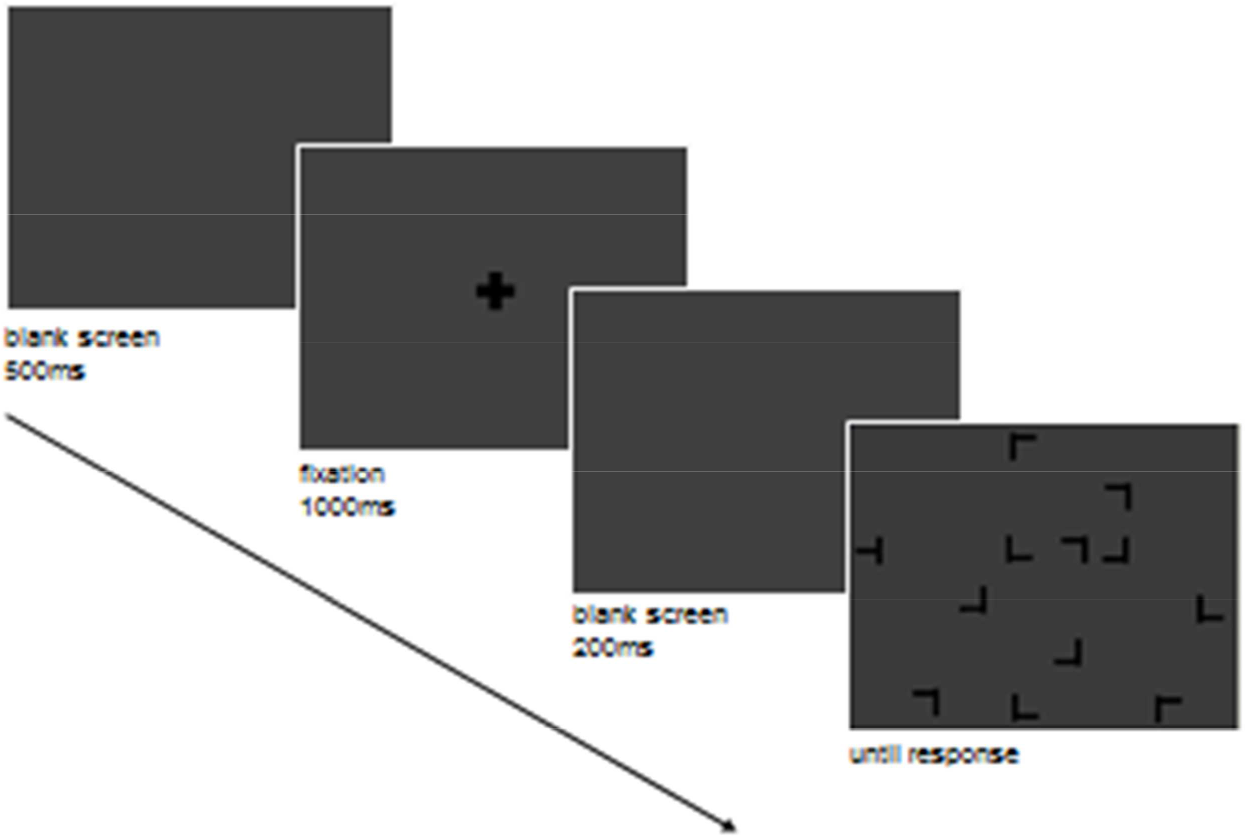
Procedure of a trial in Experiment 2. A trial consisted of the presentation of a blank screen [500ms], a fixation cross [1000ms], a blank screen [200ms], and the search display [presented until response].

### Procedures

Subjects were asked to search for the target letter T among L-shaped distractors and further to specify as quickly and accurately as possible whether the stem of the T was pointing to the left or right by mouse button presses. Each participant completed a training block, one experimental session (20 blocks), and a recognition test. The procedure of Experiment 2 was otherwise identical to the first experiment (see Figure 4).

### Data Exclusion Criteria

The same data exclusion criteria were applied for the analysis of reaction times as in Experiment 1. The data exclusion procedure led to the rejection of 3.0% (*SD* = 1.6%) of invalid data for the control group, 3.4% (*SD* = 1.5%) for handball players, and 3.4% (*SD* = 2.4%) for action video game players.

## Results

### Search Times

As in Experiment 1, blocks were aggregated into four epochs, each containing five blocks, for the search time analysis (Figure 5). A repeated measures ANOVA with configuration (repeated, novel) and epoch (1, 4) as within-subjects factors and group (control, handball, video) as between subjects factor was performed on search times. We observed significant main effects of epoch, *F*(1,72) = 265.934, *p* < 0.05, and configuration, *F*(1,72) = 47.738, *p* < 0.05. Search times decreased over time and were faster in the repeated displays (1473 ms) than in the novel displays (1565 ms). In contrast to Experiment 1, the main effect of group was not significant, *F*(2,72) = 1. 249, *p* > 0.05, indicating that overall search speed was comparable between handball players (1517 ms), action video game players (1449 ms), and controls (1591 ms). A significant interaction between epoch and configuration, *F*(1,72) = 12.872, *p* < 0.05, again reflects the increasing advantage for repeated displays over the course of the experiment. The other interactions were not significant (all *F* < 2.78, *p* > 0.05), revealing that the development as well as the overall magnitude of contextual cueing was comparable between groups.

**Figure 5.**
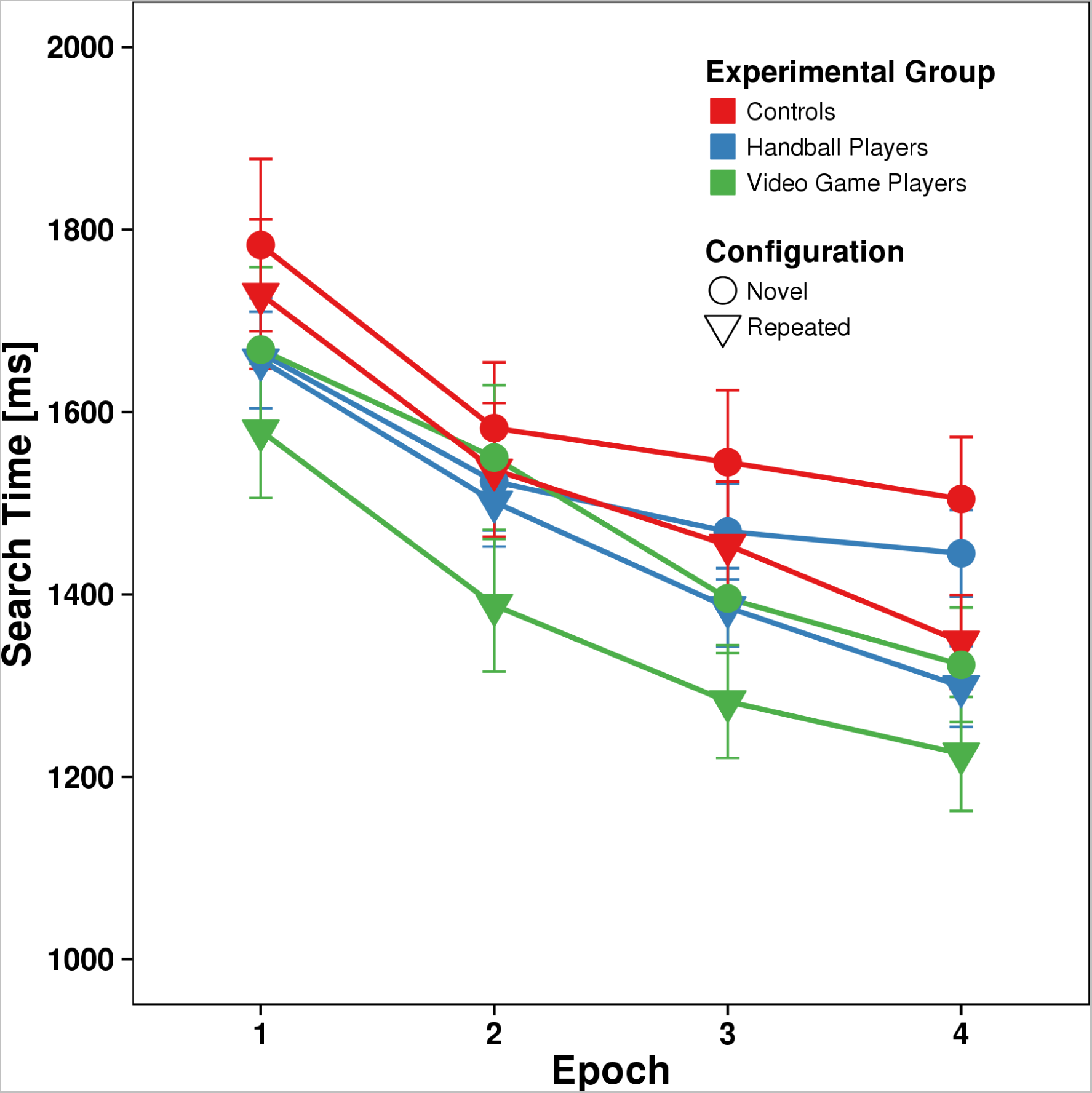
Search performance in Experiment 2. Averaged search 893 times in the visual search task for the three groups controls (red), handball players (blue), and video game players (green) for repeated (triangles) and novel (circles) display as a function of epoch. Error bars represent the standard error of the mean.

In order to investigate if the variation of the search display had a specific effect on the handball players, we ran an ANOVA on search times in Epoch 4 - after sufficient familiarization with the respective display type - across Experiments 1 and 2, with Experiment (1, 2) and group (handball, video) as between-subjects factors and configuration (repeated, novel) as within-subjects factor. Ten participants (3 handball players, 7 action video game players) who already participated in the first experiment were excluded in order to eliminate general learning effects across experiments. We observed significant main effects of experiment (Experiment 1 = 1102 ms, Experiment 2 = 1353 ms), *F*(1,84) = 29.018, *p* < 0.05, and configuration, *F*(1,84) = 49.176, *p* < 0.05, the latter again revealing that search in the repeated displays (1174 ms) was faster than search in novel displays (1258 ms). In addition, a significant interaction of experiment × configuration, *F*(1,84) = 7.884, *p* < 0.05, was found. The other interactions were not significant (all *F* < 0.71, *p* > 0.05). In particular, the non-significant interaction between experiment × group, F(1,84) = 0.489, p > 0.05, revealed that the variation of the search display had no effect on the handball players.

### Age Differences

The same age analysis as in Experiment 1 was applied to assess the effect of expertise level. With respect to age differences in the handball group, no group or interaction effects were significant (all F < 1.39, p > 0.05), indicating that both groups (junior and adult) did not differ in contextual cueing.

### Accuracy

The accuracy of the control group ranged from 94.6 to 100.0% (average 99.2%). Regarding the test groups, the accuracies varied from 95.0 to 100.0% (average 98.5%) for handball players and from 87.5 to 100.0% (average 98.6%) for action video game players. A repeated measures ANOVA on errors with the between-subject factor group (control, handball, video) and the within-subject factor configuration (repeated vs. novel) revealed neither significant main effects of group, *F*(2,72) = 1.017, *p* > 0.05, nor configuration, *F*(1,72) = 0.018, *p* > 0.05, nor a significant interaction between group and condition, *F*(2,72) = 0.162, *p* > 0.05.

### Recognition Task

Recognition analyses were identical to Experiment 1. The control group reached a mean recognition accuracy of 52.7% (*SD* = 9.5%). The mean hit rate (49.7%, *SD* = 15.7%) and the mean false alarm rate (44.3%, *SD* = 17.1%) did not differ significantly, *t*(24) = 1.398, *p* > 0.05. Recognition accuracy did not correlate with the normalized contextual cueing effect (*r* = −0.057, p > 0.05).

Mean recognition accuracy for the handball players was 52.7% (*SD* = 9.5%) with a mean hit rate of 53% (*SD* = 11.8%) and a mean false alarm rate of 40.7% (*SD* = 15.5%). Hit and false alarm rates were significantly different, *t*(24) = 3.201, *p* < 0.05, suggesting explicit learning. The normalized contextual cueing effect, however, did not correlate with recognition accuracy (*r* = −0.137, *p* > 0.05).

Action video game players obtained a mean recognition accuracy of 59.8% (*SD* = 11.4%). Participants properly categorized repeated displays as repeated on 60.3% (*SD* = 14.9%) of trials and wrongly classified 40.7% (*SD* = 19.3%) of novel displays as repeated. The difference of hit rate and false alarm rate was significant, *t*(24) = 4.312, *p* < 0.05. However, the correlation between the standardized contextual cueing effect and the recognition accuracy was not significant (*r* = 0.206, *p* > 0.05).

## Discussion

Experiment 2 yielded the typical contextual cueing pattern, with faster search in repeated displays and an increasing search advantage for repeated displays over time across groups.

Thus, contextual cueing was observed in a standard T-among-L search task, in line with the literature on subjects without particular proficiency in sports or action video game playing. Overall, the groups did not differ in the amount of contextual cueing, measured as the search time advantage for repeated over novel displays. The handball players, whose search times in Experiment 1 were faster than those of the control group, lost this advantage in Experiment 2. This is most likely due to the lack of sport-specific displays in Experiment 2. However, our data revealed that display variation had no specific effect on handball players compared to action video game players and controls.

## General Discussion

We have investigated spatial contextual cueing of visual search in expert handball players and action video game players. In Experiment 1, the search displays resembled a handball field, whereas in Experiment 2, a symbolic T-among-L search was utilized. In both experiments, we observed the typical pattern of a general reduction of search times and faster search times for repeated than novel displays, with the latter search time advantage increasing over the course of both experiments. Thus, visual search benefitted from contextual cueing, enabling efficient memory-guided search in repeated displays. However, we found no indication of superior contextual cueing in handball players and action video game players.

The lack of a contextual cueing advantage contrasted with overall shorter search times of the handball players - in repeated and novel displays alike - in the sport-specific displays, indicating that team sport expertise does not improve basic cognitive skills. The superior search performance might result from their training in handball rather than from central differences in basic cognitive abilities or any selection effect (see Memmert et al., 2009).

The selectively faster search times of the handball players for search in Experiment 1 suggests that we were at least partly successful in creating sport-specific displays. However, our displays did not resemble real configurations of players that occur in a handball game. Thus, we cannot rule out that handball players might have shown improved contextual cueing compared with controls in more realistic handball scenes (e.g. scenes from a real handball game).

These findings support the narrow transfer hypothesis, suggesting that the knowledge structures of experts operate as conjunctions between encoded information and retrieved cues in long-term memory (Furley & Memmert, 2011). Expert performers have to restore the encoding situations in order to recover the encoded information. In this way expert performers are able to fall back on long-term working memory in their particular domain of expertise, facilitating them to adapt to situational demands of their specific performance setting. However, those domain specific abilities are not transferable to other general cognitive areas (e.g. Eccles, 2006; Furley & Memmert, 2011). In fact, several studies in the field of sport (e.g. Mann et al., 2007; Memmert, Simons, & Grimme, 2009) showed that expert benefits often disappear in non-specific settings, suggesting training effects rather than superior sensory or attentional capacity as a cause for the performance benefits of experts. For example, the use of two- and pseudo three-dimensional static displays in the present study may not appropriately reflect the dynamic character of sport due to a limited viewing angle and/or missing peripheral information, which are required for adequate decision making. The use of virtual reality may improve the ecological validity of stimulus presentations, granting more stimulus control than static displays (see Tenenbaum & Eklund, 2007). Mann and colleagues (2007) found evidence that field-based studies have produced the greatest expert-novices differences. Additionally, the response – a button press – that is not a required skill for handball playing may have attenuated or even eliminated a sport-expert advantage in that handball players were not able to adequately link stimulus characteristics to response selection (Mann et al., 2007). Handball players have a different perception-action linkage in an actual game situation, when reacting to the action of another handball player, than in an experiment where the athlete has to look at static displays of a pitch and responds with button presses. Indeed, several studies suggest that the ecological validity of an experiment influences expert performance. For example, Oudejans, Michaels, and Bakker (1997) reported superior expert performance in baseball catching decisions only when real catching was required. Real game situations may have a higher ecological validity to reproduce the expert advantage (Mann et al., 2007). Indeed, Thomas, Gallagher, and Lowry (2003) reported in a meta-analytic review that a high ecological validity of the experimental setting is accompanied by larger effect sizes. The more ecologically valid the stimulus presentation, the higher may be expert-novices differences (Tenenbaum & Eklund, 2007).

In addition, effects of expertise in sport may be also modulated by different methodological factors such as sport type, level of expertise, and sex (Nougier, Ripoll, & Stein, 1989). We had a sex imbalance between groups, mainly due to the male dominance among the action video game players which may potentially influence the results. However, regarding the effect of gender differences, different studies reveal that sport expertise reduces traditional gender effects (Alves, Voss, Boot, Deslandes, Cossich, Salles, & Kramer, 2013; Lum, Enns, & Pratt, 2002). Likewise, gender differences could be nearly removed by video game training (Feng et al., 2007).

Regarding the level of expertise, the majority of participants of our study were junior athletes. Thus, it might be argued that their level of expertise was not sufficient to detect potential differences in contextual cueing compared to non-athletes. However, Alves et al. (2013) reported that junior volleyball athletes performed similarly to the adult volleyball athletes on different performance measures. Likewise, the adult athletes playing at a regional level (3rd handball league) are probably less skilled than athletes performing at a national level and above. Nevertheless, the adult athletes in the current study fulfill the ten years rule of Simon and Chase (1973). Research on expertise in sport reveal that ten years or 10.000 hours of experience and practice in a certain domain are required to develop expert performance (see Ericsson et al. (2006), for a recent review). Findings in the present study indicate that both groups (youth and adult handball players) do not differ in the contextual cueing task.

In contrast, action video game players had overall shorter search times than controls^1^ in accordance with previous research (Dye, Green, & Bavelier, 2009b). Improved hand-eye coordination in a computer setting is probably not sufficient to explain the RT advantage of video game players (Chisholm et al., 2010). Further contributing factors may be improved allocation of spatial attention (Feng et al., 2007; Green & Bavelier, 2003, 2006a) or superior visuospatial resolution (Green & Bavelier, 2007; Greenfield, DeWinstanley, Kilpatrick, & Kaye, 1994). Training studies suggested a causal relationship between action video game experience and enhanced visual and cognitive performance (see Green & Bavelier, 2003; Li, Polat, Makous, & Bavelier, 2009) due to enhanced attentional processes and the management of resources (Green & Bavelier, 2012). The greater capacity of attention for action video game players, however, is still debated (see Irons, Remington, & McLean, 2011).

It has been pointed out that the superior performance of experts may either be due to training or, alternatively, due to selection. Subjects with superior attentional skills may find it more rewarding to play handball or action video games because they are more successful (Kristjansson, 2013). In a study by Boot et al. (2008), e.g., superior performance by expert action video game players could be replicated on a wide range of attentional and cognitive tasks. However, non-gamers did not reach superior performance even after intensive training.

The present findings, however, do not indicate that either handball training or action video game play lead to generalized contextual cueing. Nor do our data suggest a selection effect in the sense that subjects with faster memory-guided visual search may be more likely to become expert action video game players.

Thus, findings in the present study do not suggest that contextual cueing is an important limiting basic ability in the domain of team sports and action video game playing.

## Acknowledgements

This work was supported by Florian Baumgartner and Daniel Kottke by helpful statistical advice and discussions.

We would also like to thank the Sportclub Magdeburg for the opportunity to study elite athletes, the coaches for their support, and the athletes for their willingness to participate in the study.

## Footnotes

1 As a significant effect in Experiment 1 and a tendency in Experiment 2

